# Dynamics of Saccade Trajectory Modulation by Distractors: Neural Activity Patterns in the Frontal Eye Field

**DOI:** 10.1101/2024.04.04.587872

**Authors:** H. Ramezanpour, D. H. Kehoe, J. D. Schall, M. Fallah

## Abstract

The sudden appearance of a visual distractor shortly before saccade initiation can capture spatial attention and modulate the saccade trajectory in spite of the ongoing execution of the initial plan to shift gaze straight to the saccade target. To elucidate the neural correlates underlying these curved saccades, we recorded from single neurons in the frontal eye field (FEF) of two rhesus monkeys shifting gaze to a target while an isoeccentric distractor appeared either left or right of the target at various delays after target presentation. We found that the population level of pre-saccadic activity encoded the direction of the saccade trajectory. Stronger activity occurred when saccades curved toward the distractor, and weaker when saccades curved away. This relationship held whether the distractor was ipsilateral or contralateral to the recorded neurons. Meanwhile, visually responsive neurons showed asymmetrical patterns of excitatory responses that varied with the location of the distractor and the duration of distractor processing relating to attentional capture and distractor inhibition. During earlier distractor processing, neurons encoded curvature towards the distractor. During later distractor processing, neurons encoded curvature away from the distractor. This was observed when saccades curved away from distractors contralateral to the recording site and when saccades curved towards distractors ipsilateral to the recording site. These findings indicate that saccadic motor planning involves dynamic push-pull hemispheric interactions producing attraction or repulsion for potential but unselected saccade targets.

**Significant Statement:** This study not only advances our understanding of oculomotor function in dynamic environments but also has potential clinical relevance for diagnosing and understanding disorders characterized by abnormal saccade trajectories. Our research elucidates the neural mechanisms behind saccade trajectories that are not always linear due to the brain’s integration of multiple visual cues and/or motor plans. By analyzing the frontal eye field (FEF) activity in rhesus monkeys, we found that saccade directionality and timing are influenced by the interaction between FEF visual neurons representing target and distractor stimuli. The FEF’s role extends beyond a winner-takes-all approach, incorporating saccade vector averaging computations that produce curved saccades. Furthermore, ipsilateral visual neurons encode distractor suppression that drives curvature away from the distractor.

## Introduction

Saccadic target selection involves a complex neural network that integrates visual information with cognitive variables (Schall, 1995). In monkeys, the superior colliculus (SC), frontal eye fields (FEF), and lateral intraparietal area (LIP), as the three main nodes of the oculomotor system, play key roles in this process (Munoz and Everling, 2004; Bisley and Goldberg, 2010; Schall, 2015). Visual stimuli are processed to determine their relevance and priority, while cognitive goals and current tasks influence the selection by biasing attention towards certain objects or locations. The oculomotor system encodes potential targets and decides where the eyes should move next (Fecteau and Munoz, 2006).

When a sudden distractor appears just before the initiation of a saccade, it introduces a competitive element to the saccadic target selection process in the oculomotor system (Theeuwes et al., 1999; Theeuwes, 2010). The appearance of a new stimulus creates a conflict in the oculomotor system which is already engaged with the original target. Neural circuits involved in saccade programming, such as the SC (Horwitz and Newsome, 1999; Krauzlis and Dill, 2002; Krauzlis et al., 2004) and the FEF (Bruce and Goldberg, 1985; Schall and Hanes, 1993; Schall, 1995; Tehovnik et al., 2000), must rapidly assess this new information. The competition between the distractor and the planned saccade target can lead to alterations in saccade latency, trajectory, and destination (Walker et al., 1997, 2006; Van der Stigchel et al., 2007; Walker and McSorley, 2008; Jonikaitis and Belopolsky, 2014; Wollenberg et al., 2018; Castellotti et al., 2023). In some cases, if the distractor is highly salient or relevant, it might even override the original saccade plan, redirecting the eye movement to the new target (Theeuwes et al., 1999).

When a distractor appears, it can create an inhibitory or excitatory gradient within the oculomotor system (McPeek et al., 2003; Port and Wurtz, 2003; McPeek, 2006; White et al., 2012; Kehoe and Fallah, 2017), where neurons map out the spatial landscape for potential saccade targets (Fecteau and Munoz, 2006). If the brain decides to prioritize the initial target, the saccade might still show a curvature toward or away from the distractor, where the trajectory represents a vector sum of the activity evoked by the target and the distractor (Port and Wurtz, 2003). The temporal dynamics and direction of curvature depends on various factors (Van der Stigchel, 2010) such as target and distractor features (Kehoe et al., 2023), and the relative timing between the appearance of the distractor and initiation of the already planned saccade (Kehoe and Fallah, 2017; Giuricich et al., 2023). Several electrophysiological and inactivation studies have shown that saccade curvatures toward and away from distractors are correlated with higher excitation (McPeek et al., 2003; Port and Wurtz, 2003) and inhibition (Aizawa and Wurtz, 1998) in the SC neurons representing the distractor location.

Neural correlates of curved saccades in the highest order area of the oculomotor system i.e. the FEF, has been much less studied. In the only electrophysiological study specifically investigating curved saccades, it was found that an increase in perisaccadic activity of FEF neurons coding the distractor location would lead to saccades that curve toward a distractor and a decrease in perisaccadic activity would lead to saccades that curve away from the distractor (McPeek, 2006). Yet, this study did not investigate the temporal dynamics of target-distractor competitions which result in modulations of saccade trajectories. To better understand these neural mechanisms, we conducted an experiment in which monkeys had to make saccades to a visual target in the presence of a competing distractor that was presented at various time points relative to target onset while we were recording from single units in the FEF. We analyzed the neuronal responses in relation to distractor processing time, i.e. from distractor onset to saccade onset, as this is the period of target-distractor competition that dynamically determines the curvature of a saccade.

## Materials and Methods

Two male macaques underwent a surgical procedure to implant a headpost and a recording cylinder over the right (monkey D) and left (monkey K) FEF. The recording chamber was centered in stereotaxic coordinates at 25 mm anterior for both monkeys, and 19 and 20 mm lateral for monkeys D and K, respectively. The recorded sites were confirmed to be within the low-threshold FEF (<50 µA) using microstimulation criteria defined by Bruce and Goldberg (1985). All experimental and surgical procedures complied with animal care guidelines, as defined by the CCAC (Canadian Council on Animal Care). The study and all associated protocols were approved by York University’s Animal Care Committee.

### Apparatus and Measurement

Experimental control was maintained using Cortex software (http://dally.nimh.nih.gov/). Eye gaze was tracked using an infrared eye tracker (ISCAN model ETL-200, 240 Hz). Stimuli were presented on a computer monitor (Viewsonic G225f, 1,024 × 768 resolution, 60 Hz) that was placed 50 cm from the monkey. For neural recordings, a precision microdrive (Crist Instruments) was employed to position a tungsten microelectrode (FHC) accurately. Neural activity was monitored in real-time and recorded using a high-fidelity neural data acquisition system (Plexon). Neurons were isolated offline using Offline Sorter (Plexon) for subsequent analyses.

### Visual Stimuli and Experimental Task

The saccade target was a white circle (luminance = 122.70 cd/m^2^) that subtended 0.3° in diameter and was located 10° above the central fixation. The distractor had the same size and luminance as the target. Stimuli were embedded in a black background. Trials were started when the monkeys held their gaze on a white central fixation point (0.3° diameter) set within a 3° square boundary for a duration of 500 ms. Following this half-second period, the fixation point disappeared and simultaneously a target appeared 10° above the central fixation (see **Figure 1**). The monkeys were then required to direct their gaze to this newly presented target immediately upon its appearance. The distractor was introduced after a variable delay of 16, 50, 100, 150, or 200 ms at 10° eccentricity and at either +/- 40° angle from the target, this timing delay being designated as the distractor target onset asynchrony (DTOA). The completion of a trial was marked by either a saccadic movement toward the target or the lapse of 500 ms (signaling a time-out). An intertrial interval (ITI) of one second with a dark screen was used between trials. The position of the distractor and the DTOA were varied randomly in each trial.

**Figure 1.**
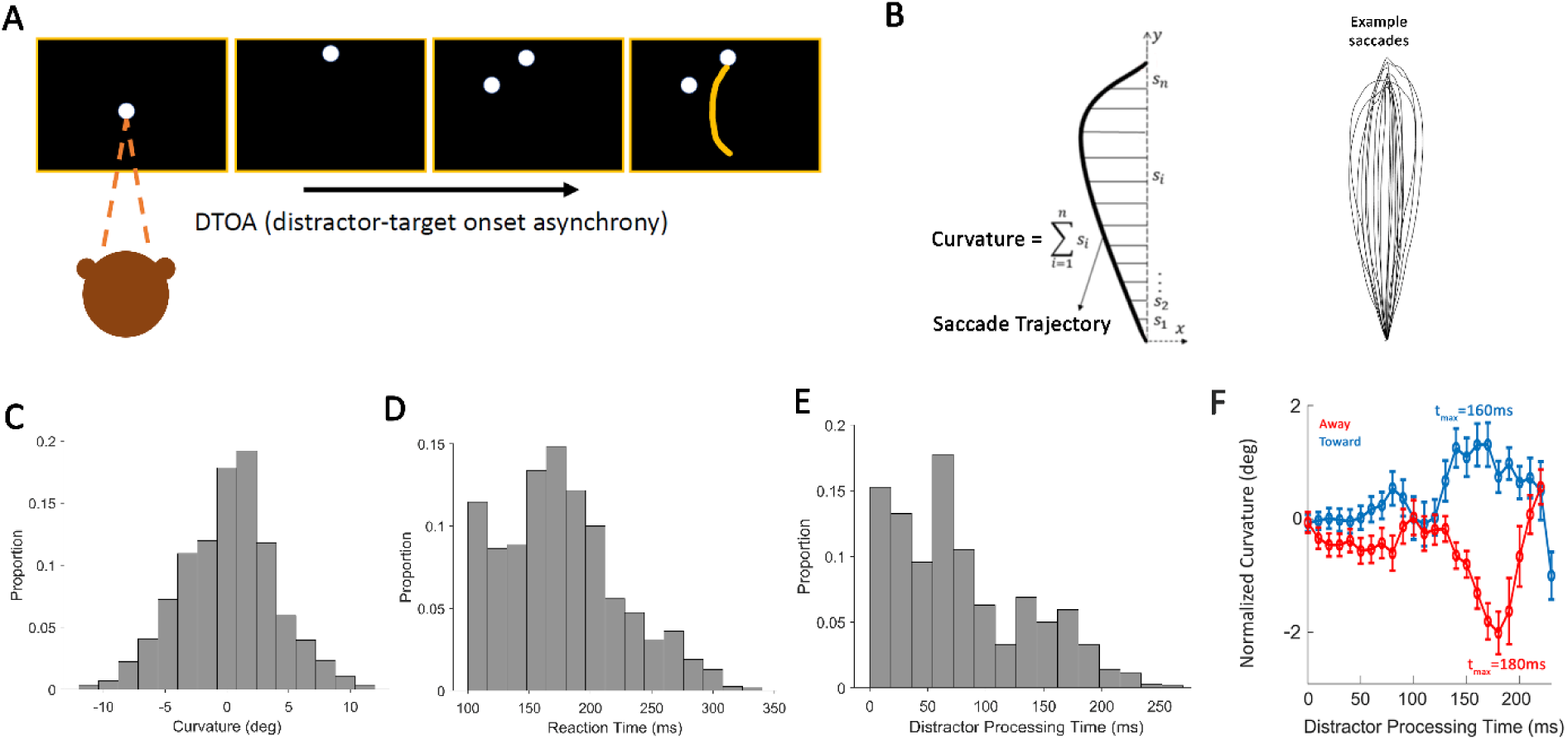
(A) Schematic of the experimental paradigm. A trial starts with the monkey fixating on a central point before moving his gaze to a target. Followed by the target presentation a distractor appears at different intervals (DTOA) either on the left or right side of the target. The monkey has to ignore the distractor and make a vertical saccade to the target. While the monkey is performing this task single units are recorded from his FEF. (B) Saccade curvature calculations. The saccade trajectory is broken down into vectors to quantify the deviation from a straight line (Kehoe and Fallah, 2017; Kehoe et al., 2018). (C-E) are histograms representing the distribution of saccade curvature, saccadic reaction times, and distractor processing times, respectively, showcasing the variability and range of each metric. (F) Normalized curvature of saccadic trajectories away from (red) and towards (blue) a distractor over time, with peak deviation times (t_max_) indicating the maximum influence of the distractor. Curvature values for toward and away are significantly different in the window 140 – 200 ms after distractor onset (Wilcoxon sign-rank test, p<0.05).

### Saccade Detection

Trials that contained blinks were excluded from further analysis. Saccades were defined based on a velocity threshold of 20°/s for at least 8 ms and a peak velocity exceeding 50°/s. Saccades were excluded from further analysis if they were accompanied by more than one corrective saccade or one corrective saccade larger than 2° visual angle. Also, saccades whose amplitude was smaller than 2° visual angle or saccades with less than 100 ms latency were excluded. Other exclusion criteria were endpoint deviations larger than 2° and pre-saccadic drift larger than 2°. To assess the trajectory of saccadic eye movements, the initial points of the saccades were repositioned to a common origin and then mathematically rotated to align the final points along the positive Y-axis. The latency of the saccadic response, or saccade reaction time, was determined by the interval from the appearance of the target to the commencement of the saccade. The curvature was quantified by tallying all the perpendicular deviations from an imaginary direct line that spans from the beginning to the end of the saccade path (as depicted in **Figure 1B**). Twenty exemplary saccade trajectories have been plotted in **Figure 1B**. To accommodate for the inherent curvature present in natural saccadic movements, we established a baseline curvature for each monkey by measuring saccades directed at the same target without any distracting stimuli. This baseline curvature was then deducted from the curvature measurements of all saccades to normalize for the innate curvature characteristics of the saccades. Saccade curvatures were analyzed as a function of distractor processing time, i.e. the interval between distractor onset to saccade onset. For further analysis, we median-split the saccade curvature distribution to high- and low-curvature values. These values were used to investigate whether low- and high-curvature saccades are differentially processed by FEF neurons.

### Statistical Analysis and Functional Categorization of FEF Neurons

Statistical comparisons were performed using non-parametric tests (Wilcoxon sign-rank test, rank-sum test). To obtain the time windows in which the firing rates between two conditions were statistically different, the p<0.05 threshold had to be met consecutively for all bins for at least 10 ms to avoid multiple comparison problems. To categorize neurons as visual, visuomotor, and motor neurons, the definitions from (Bruce and Goldberg, 1985) were adopted. We compared the motor response (in the 50 ms before saccade onset) and the visual response (50–150 ms after target onset) with a baseline response (100 ms before target onset) from that trial. We used Wilcoxon sign-rank test at the p < 0.05 level to indicate significance. Out of 104 recorded FEF neurons, 93 neurons were task related, i.e. showed a significant response in one of these windows, ensuring that either the target or the distractor was overlapping with the receptive field of the neuron. Neurons were categorized as motor neurons if only the motor response was significantly higher than the baseline response. Neurons were categorized as visual neurons if only the visual response was significantly higher than the baseline response. Neurons were categorized as visuomotor neurons if both the visual and motor responses were significantly higher than the baseline response. Since there is a stronger representation of the contralateral visual field in the FEF, we investigated the relationship between low- and high-curvature saccades and FEF neural activity for distractors inside the contra- or ipsilateral field separately. All of the analyses were performed in Matlab.

## Results

Figure 1 presents three histograms (C, D, E), each illustrating different aspects of saccadic eye movements toward a target in the presence of a visual distractor. Panel C shows the distribution of saccade curvatures (negative values indicating curvature away from the distractor, and positive values toward the target). Note that the symmetrical distribution around 0 degrees suggests a roughly equal propensity for saccades to curve toward or away from the target in the presence of distractors, across the range of DTOAs tested. Panel D depicts the overall distribution of reaction times. The distribution is skewed to the left, indicating that most of the saccades have a short reaction time, with fewer saccades taking longer to initiate. This was largely due to the fact that the two monkeys were performing this task for several months, and they were overtrained with respect to the task and structure of trials. Note that anticipatory saccades (reaction times less than 100 ms) were excluded from analysis. Panel E provides the distribution of distractor processing times (reaction time - DTOA). Figure 1F depicts curvature modulation as a function of distractor processing time. Curvature values for saccades curved toward and away from the distractor (positive and negative values) were normalized by setting the mean of initial curvature values (distractor processing time =0) to zero. The data points illustrate a bimodal peak in curvature deviation. Although, the initial modulations for both toward and away conditions (distractor processing time less than 100 ms) were almost identical, saccades curving towards the distractor peaked at 160 ms and those curving away peaked at 180 ms after the distractor presentation.

To investigate how FEF neural responses encode saccade curvature, we first calculated the population activity of all task-related FEF neurons immediately prior to saccade initiation. Figure 2A demonstrates the population firing rate when saccades are curved toward the distractor on the contralateral side, representing attentional capture by the distractor. As compared to saccades with lower curvature, there’s a marked increase in the firing rate for saccades with larger curvature when saccades curved toward the contralateral distractor in a window of 35–90 ms before saccade onset. Figure 2B illustrates the firing rate when saccades are curved away from the distractor on the contralateral side, representing distractor suppression, exhibiting a different pattern than Panel A, with a larger peak firing rate for saccades with lower curvature in a window of 46–72 ms before saccade onset. There was a similar pattern for the ipsilateral distractor i.e., higher curvature was associated with a higher firing rate when saccades were curved toward the distractor in a window of 20–111 ms prior to saccade onset (Figure 2C), and reversely, lower curvature was associated with a higher firing rate when saccades curved away from the ipsilateral distractor in a window of 37–141 ms (Figure 2D). In Figure 2A-D, the average saccade trajectories for the two spike density functions are displayed on the right, with low curvature in green and high curvature in black.

**Figure 2.**
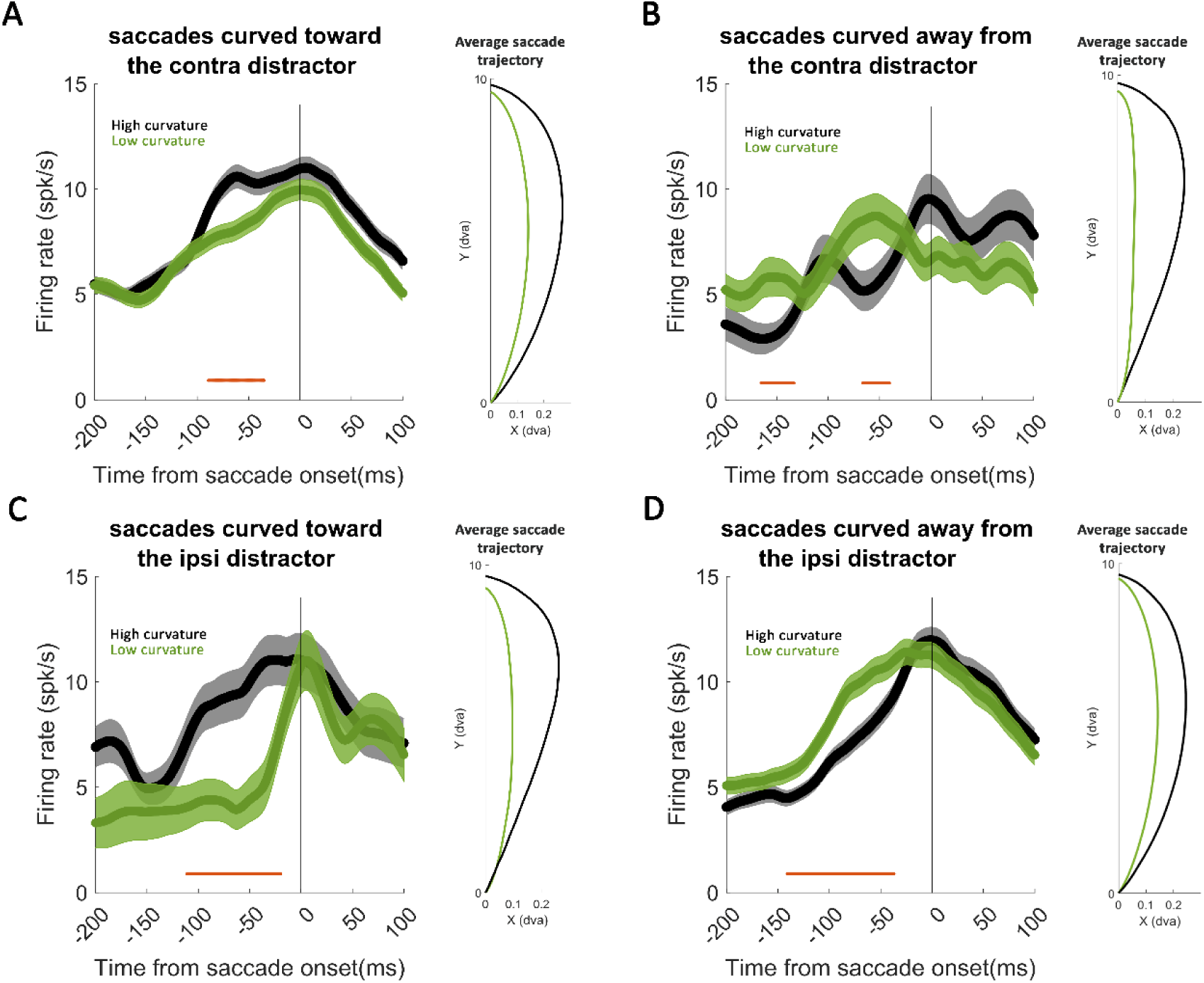
Neuronal response profiles during saccadic eye movements in the presence of a visual distractor. The graphs depict the mean firing rate of the FEF neuronal population recorded over time relative to the onset of a saccade. (A) Represents the firing rate when saccades curved toward the contralateral distractor, showing an elevated response in a window of 35–90 ms just before saccade onset for more curved saccades. The average saccade trajectory for the two spike density functions is displayed on the right. (B) Illustrates the firing rate when saccades are curved away from the distractor on the contralateral side. There was a stronger firing to saccades with lower curvature in windows of 46–72 ms, and 139–171 ms before saccade onset. (C-D) Firing patterns for ipsilateral distractors are very similar to contralateral ones with slight timing differences. Higher curvature was associated with a higher firing rate when saccades were curved toward the distractor in a window of 20–111 ms prior to saccade onset (C), and reversely, lower curvature was associated with a higher firing rate when saccades curved away from the ipsilateral distractor in a window of 37–141 ms (D). Shaded areas represent the standard error of the mean. The horizontal red bars indicate the duration of the significant differences in firing rates. The vertical line at time 0 indicates the onset of the saccade. All statistical comparison were performed using Wilcoxon sign-rank test, and a p<0.05 threshold holding for at least 20ms. High and low curvature trials are based by median splitting.

In order to further investigate the relationship between saccade curvature and FEF neural activity, we then separated the FEF neurons into visual (n=21), motor (n=5) and visuomotor (n=67) (see method section). Because the number of motor neurons in our sample was too small for further analysis, we only investigated the time course of distractor processing in the visual and visuomotor neurons. Among the sample of visually responsive neurons in the FEF, we found unbalanced patterns of excitatory responses before saccades curved toward or away from the distractor. This varied with distractor location and processing time. For contralateral distractors (Figure 3A), an excitatory visual response was associated with saccades curved toward the distractor, which emerged earlier than the response for saccades curved away from the distractor (toward: time of peak =100 ms, latency=48 ms; away: time of peak = 166ms, latency=140 ms). Latencies were calculated as the first time point after distractor onset for which the activity was significantly higher than the 50 ms baseline activity immediately prior to distractor onset (p<0.05, Wilcoxon sign-rank test). A sliding window statistical comparison showed that excitatory visual response encoding saccades curved toward the distractor was significantly different than the response encoding saccades curved away from the distractor in two windows of 42–58 ms and 86–118 ms after distractor onset (Wilcoxon sign-rank test, p<0.05). For ipsilateral distractors (Figure 3B) this pattern was reversed—shorter-latency visual responses when related to saccades curved away from the distractor emerged earlier than visual responses when saccades curved towards a contralateral distractor (away: time of peak=101 ms, latency=72 ms; toward: time of peak = 172 ms, latency=136 ms). A sliding window statistical comparison showed that visual responses encoding saccades curved toward the distractor was significantly different than the response encoding saccades curved away from the distractor in two windows of 70–124 ms and 150–81 ms after ipsilateral distractor onset (Wilcoxon sign-rank test, p<0.05). Similar peak response time to saccades curved toward a contralateral distractor or curved away from an ipsilateral distractor (100 ms and 101 ms respectively, see Figure 3A-B), and peak response time to saccades curved away from a contralateral distractor or curved toward an ipsilateral distractor (166 ms and 172 ms respectively, see Figure 3A-B) suggest that direction of curvature is encoded in the peak activity of both contralateral and ipsilateral distractor responding FEF neurons in two different time windows, and regardless of the contra/ipsi designation of the neuron’s visual receptive field. The firing rate of visuomotor neurons lacked any specific discriminatory power between the curvature toward and away conditions split by contralateral distractors (Figure 3C) or ipsilateral distractors (Figure 3D).

**Figure 3.**
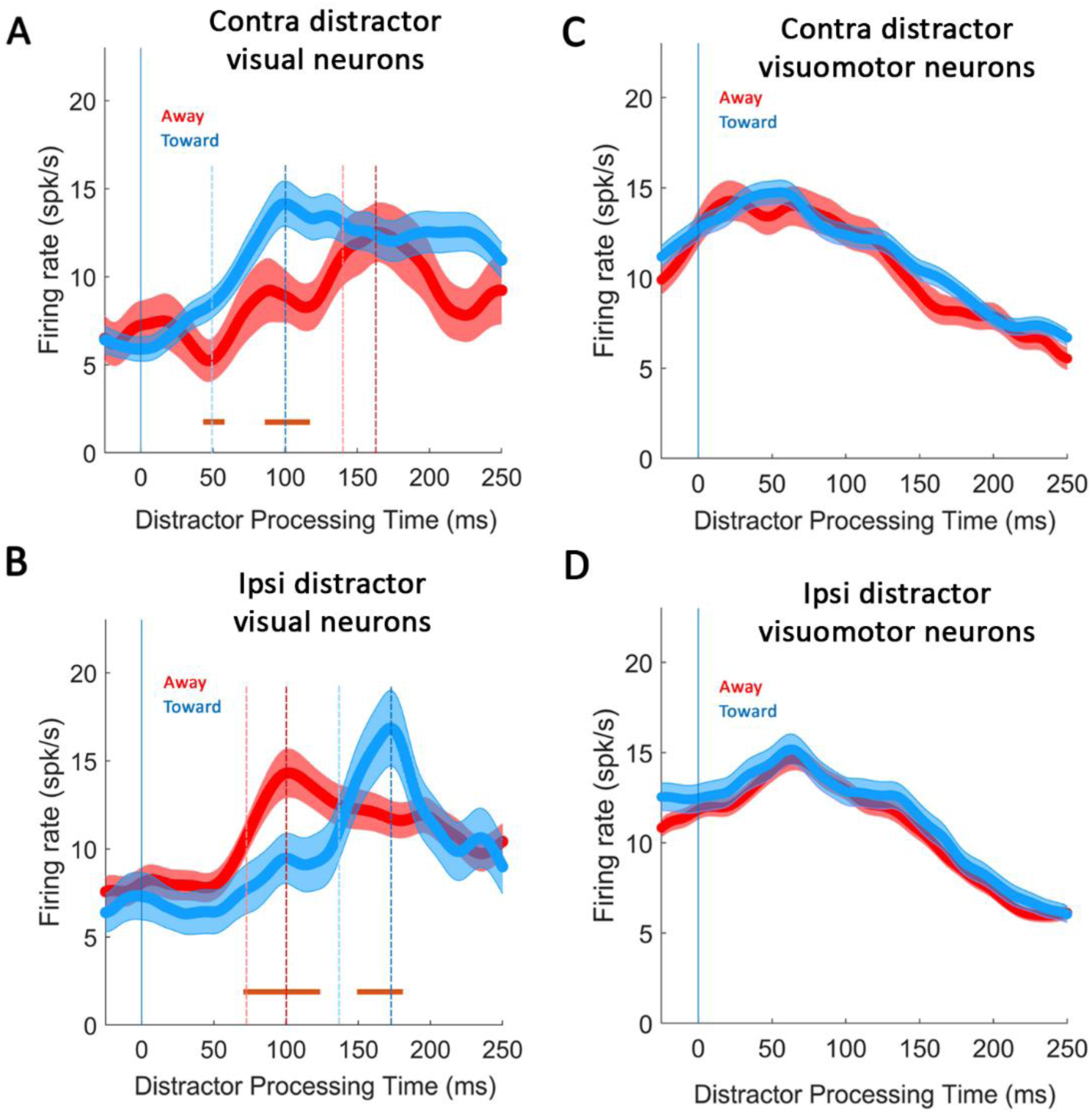
(A) Mean firing rate of visual neurons in response to a contralateral distractor, differentiated by saccades moving away (red) and towards (blue) the distractor, showing distinct temporal profiles and latency to peak firing rate (t_max_ and latency). Light blue and red dashed lines represent latency for toward and away conditions respectively, and dark blue and red dashed lines represent time of the peak firing (t_max_) for toward and away conditions respectively. (B) Neuronal firing rate responses to an ipsilateral distractor, with the same color coding for directionality as in (A), showing the timing and amplitude differences in neuronal activity. Light blue and red dashed lines represent latency for toward and away conditions respectively, and dark blue and red dashed lines represent time of the peak firing (t_max_) for toward and away conditions respectively. (C) Mean firing rate of visuomotor neurons in response to a contralateral distractor, differentiated by saccades moving away (red) and towards (blue) the distractor. (D) Mean firing rate of visuomotor neurons in response to an ipsilateral distractor. Error bars and shaded regions represent standard error of the mean. The horizontal red bars indicate the duration of significant differences between the ‘away’ and ‘towards’ conditions (Wilcoxon sign-rank test, p<0.05).

## Discussion

We recorded FEF neuronal activity while monkeys executed target-directed saccades and ignored peripheral distractors that abruptly onset at a random interval after target onset. Targets always appeared along the vertical meridian and distractors unpredictably appeared 40° isoeccentrically to either the left or right of the target.

We observed that FEF neuronal activation was greater in the perisaccadic interval for saccades curved more towards distractors than for saccades curved less, while conversely, activation was lower in the perisaccadic interval for saccades curved away more from distractors than for saccades curved away less from the distractors. This pattern of results replicates earlier work investigating the association between saccade curvature and perisaccadic neuronal activation in the FEF (McPeek, 2006) and SC (McPeek et al., 2003; White et al., 2012). Interestingly, we observed this effect regardless of whether (1) the curvature was directed towards or away from distractors in the visual- and/or motorfield of the cell and (2) the curvature was directed towards or away from distractors in the opposite hemifield. This unexpected result suggests that while saccade curvature is *driven* by spatiotemporal averaging of perisaccadic activation in oculomotor substrates (McPeek et al., 2003; Port and Wurtz, 2003; McPeek, 2006; White et al., 2012) it is also *compensated* for via opposing perisaccadic activation patterns within the same substrates. That is, for saccades curved towards a distractor in the ipsilateral hemifield relative to the recording site, perisaccadic FEF neuronal activation should pull the saccade in the opposite direction of the observed curvature. This activation can therefore only serve to compensate for the spatial biasing of the saccade. To further investigate this reasoning, we subsequently examined FEF activation as a function of time since distractor onset, split by distractor hemifield and saccade curvature direction.

We observed that saccades were maximally curved towards distractors after 160 ms of distractor processing time while saccades were maximally curved away from distractors after 180 ms of distractor processing time. This replicates the temporal onset of saccade curvature observed in humans under similar experimental conditions (Kehoe and Fallah, 2017; Kehoe et al., 2023), but shows much less differentiation between the time course of saccades curved towards or away from distractors as seen previously in humans (Kehoe and Fallah, 2017). That is, for the monkeys examined here, there was a high degree of overlap in the time course for saccades curved towards and away from distractors, which may likely be an effect of overtraining.

On early distractor processing time scales, FEF neurons encoded the movement direction of curvature. When distractors appeared in a contralateral receptive field of the visual neuron and saccades curved towards the distractor, we observed a typical visual onset response with a latency of 48 ms, and peaking at 100 ms after distractor onset (blue curve in Figure 3A). When saccades curved away from the distractor, there was a similar time to peak (101 ms) but a slightly longer latency (72 ms), but for visual neurons with ipsilateral receptive fields (red curve in Figure 3B). In both of these curvature direction conditions, FEF visual onset responses were encoding saccadic spatial biasing dependent on whether the ipsilateral or contralateral receptive field visual neurons were the active population during the peak activity around 100 ms. Previous research has shown that saccade curvature away from a distractor is associated with lower neuronal activation in oculomotor cells encoding the distractor than for corresponding straight saccades (White et al., 2012). Past theorizing on the neural correlates of saccade curvature away from distractors has argued that this firing rate may introduce a negative contribution in a vector weighted average of target and distractor vector codes in oculomotor neurons (McSorley et al., 2006, 2009; Kehoe and Fallah, 2017; Kehoe et al., 2021). However, here, we show that saccade curvature away from distractors is actually associated with a positive contribution from ipsilateral oculomotor cells encoding movement directions *opposite* of the distractor. Visual FEF neurons with ipsilateral visual receptive fields, similar to those with contralateral receptive fields, engage in ipsilateral distractor suppression leading to curvature away from the distractor likely from the planning of contralateral saccades, as evidenced by the direction of curvature, rather than in the direction of their receptive fields. It is interesting to note that curvatures towards or away from the distractor was differentiated by activity within visual neurons, but not within the larger population of visuomotor neurons.

A similar effect was first observed by (McPeek et al., 2003) for saccades curved towards distractors, but with an experimental paradigm in which distractors simultaneously appeared in both the ipsi- and contralateral visual hemifields. As such, they could not disentangle a positive contribution of directionally-congruent neuronal activation and a negative contribution of directionally-incongruent neural suppression to the spatial average computation. Here, we elucidate the underlying mechanisms for those observations by showing that saccade curvature away from distractors is at least partially driven by a positive contribution of directionally-incongruent activation, ipsilateral visual neurons.

Intriguingly, at later distractor processing time scales (red curve in Figure 3A and blue curve in Figure 3B), we observed that FEF neurons encoded the anti-movement direction of curvature. When saccades curved in the opposite direction relative to the cell’s receptive field, we saw a swell of neuronal activation with a latency of approximately 140 ms and a peak around 170 ms. However, the directional biases in curvature were already decodable from the earlier peak activity around 100 ms and whether the visual neurons had contra- or ipsi-lateral receptive fields. Thus, these peaks likely would be due to cross-hemisphere activation (See Figure 4). That is, when saccades curved towards the distractor, early activity occurred in contralateral receptive field visual neurons in the contralateral FEF, which would then cross to the ipsilateral FEF to ipsilateral RF visual neurons, producing the later peak. The reverse occurs for curvatures away from the distractor: early activity in the ipsilateral RF visual neurons in the ipsilateral FEF would then likely cross to the contralateral visual neurons in the contralateral FEF producing their later peak. These results therefore suggest that saccade trajectory deviations arise from a complex push-pull interaction of visual FEF neurons across hemispheres, and specifically provide a role for ipsilateral RF visual neurons in producing curvature away from a putatively suppressed distractor. The existence of commissural fibers between FEFs has been already established (Gould et al., 1986). This proposal has been depicted in Figure 4 which conceptualizes the interhemispheric interaction between contra- and ipsi-lateral visual neurons in both FEFs during saccade planning in the presence of a distractor. Further studies are needed to test this model. Overall, activity patterns in FEF visual neurons distinguish between saccade trajectories deviated towards or away from a distractor.

**Figure 4.**
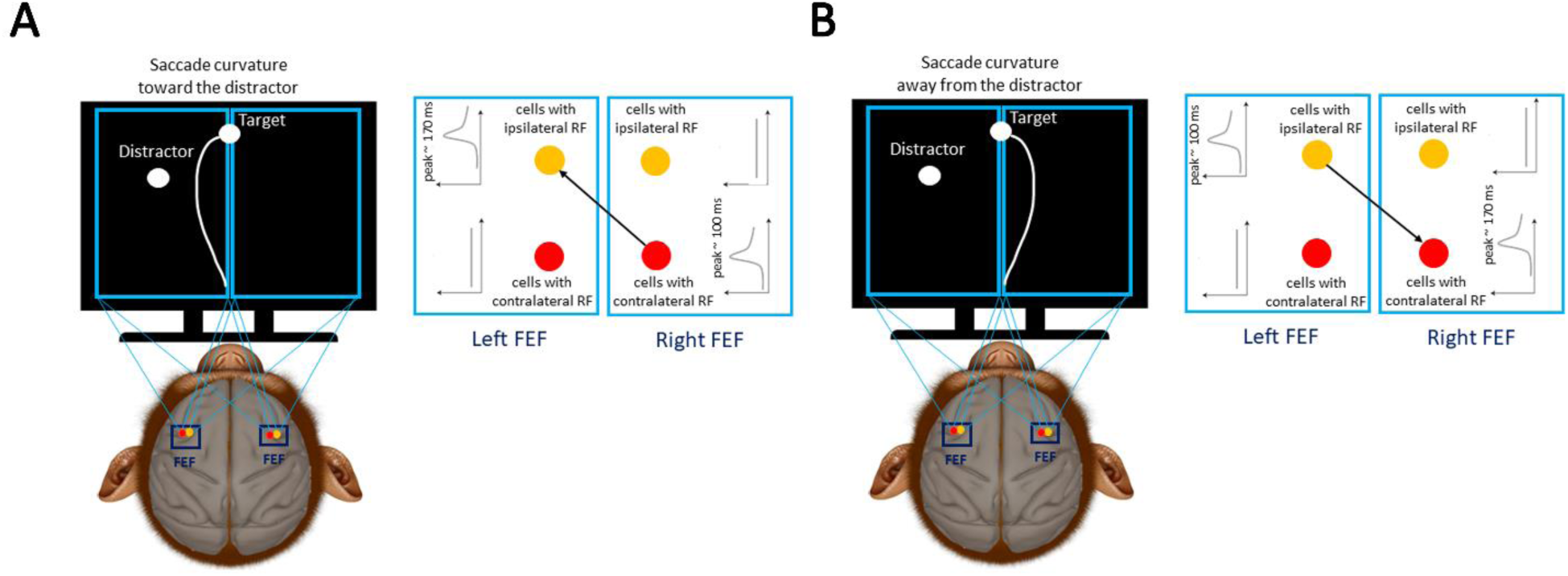
Comparative illustration of saccade trajectories, associated FEF neural population activity and cross-talk between the two hemispheres. **(A)** A representative saccade trajectory curved towards the distractor is shown. In this case, the FEF population activity of neurons with contralateral receptive fields, representing the distractor location rises with a peak at around 100 ms after distractor onset which is followed by the other hemisphere FEF neurons with ipsilateral receptive field at around 170 ms after distractor onset (black arrow). The downstream motor neurons (not shown here) representing the motor field locations get a time-revolving population vector encoding the distractor location which is vector averaged with the already planned saccade target population vector. **(B)** Similar to **(A)** except that neurons with ipsilateral receptive fields in the left FEF lead the activity of neurons with the contralateral receptive field in the right FEF leading to saccades curved away from the distractor. Note that in both A and B, there are FEF visual neurons with empty receptive fields that remain silent.

